## Supplementary figures and images for "Virological characteristics of the SARS-CoV-2 XBB.1.5 variant"

### Supplemental Figures

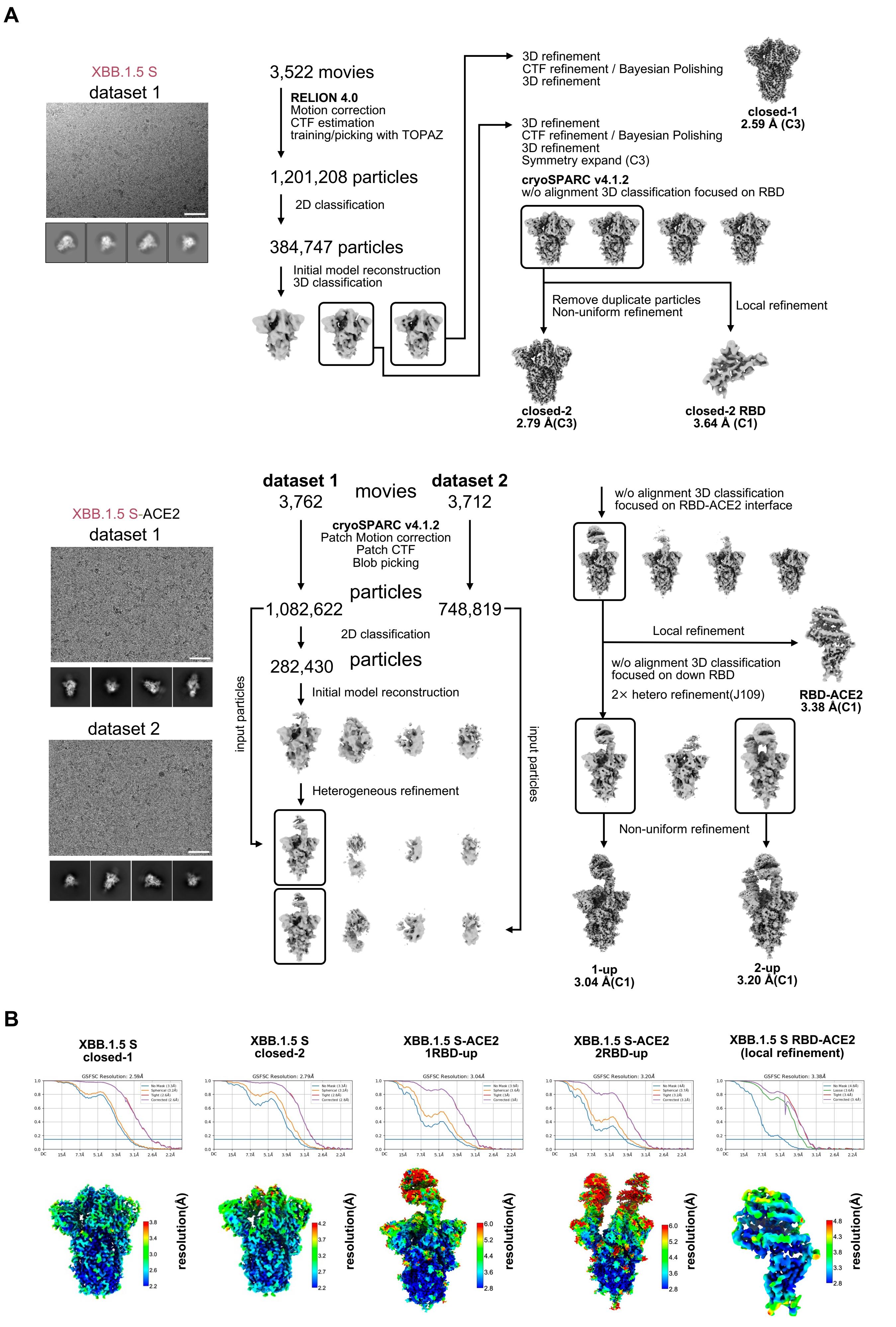

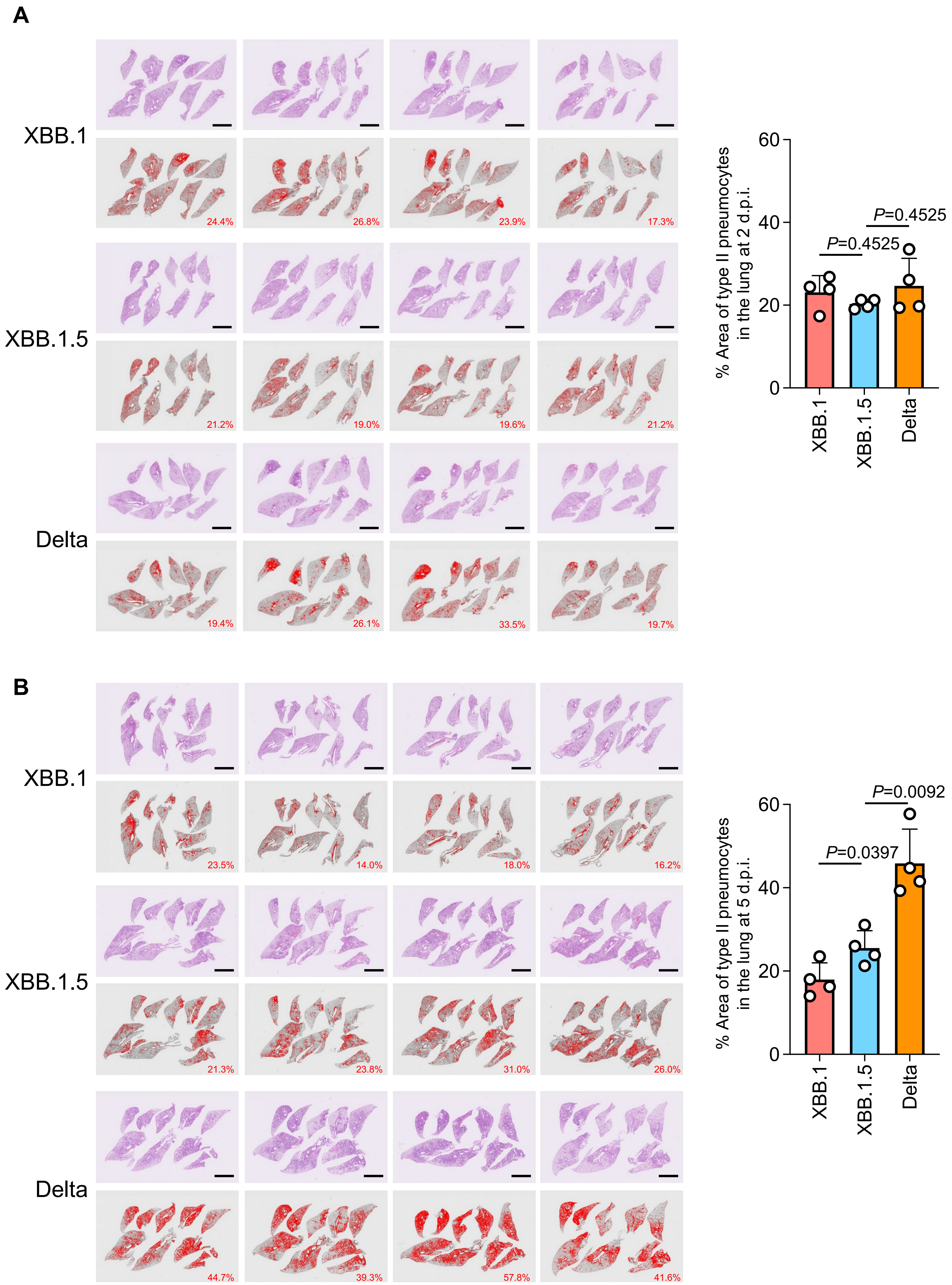

**Figure S3**

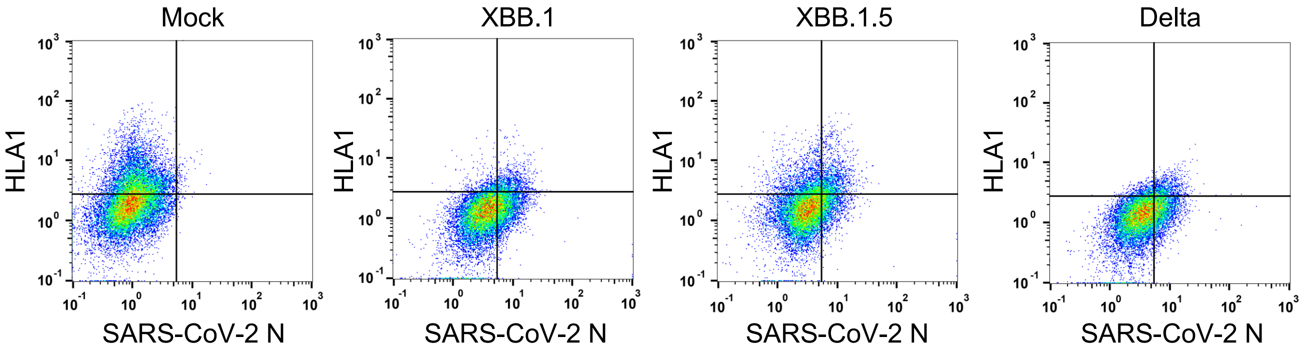
